# Polygenic Risk Score Associates with Atherosclerotic Plaque Characteristics at Autopsy

**DOI:** 10.1101/2023.07.05.547891

**Authors:** Anne Cornelissen, Neel V. Gadhoke, Kathleen Ryan, Chani J. Hodonsky, Rebecca Mitchell, Nathan Bihlmeyer, ThuyVy Duong, Zhifen Chen, Armelle Dikongue, Atsushi Sakamoto, Yu Sato, Rika Kawakami, Masayuki Mori, Kenji Kawai, Raquel Fernandez, Saikat Kumar B. Ghosh, Ryan Braumann, Biniyam Abebe, Robert Kutys, Matthew Kutyna, Maria E. Romero, Frank D. Kolodgie, Clint L. Miller, Charles C. Hong, Megan L. Grove, Jennifer A. Brody, Nona Sotoodehnia, Dan E. Arking, Heribert Schunkert, Braxton D. Mitchell, Liang Guo, Renu Virmani, Aloke V. Finn

## Abstract

**Background:** Polygenic risk scores (PRS) for coronary artery disease (CAD) potentially improve cardiovascular risk prediction. However, their relationship with histopathologic features of CAD has never been examined systematically.

**Methods:** From 4,327 subjects referred to CVPath by the State of Maryland Office Chief Medical Examiner (OCME) for sudden death between 1994 and 2015, 2,455 cases were randomly selected for genotyping. We generated PRS from 291 known CAD risk loci. Detailed histopathologic examination of the coronary arteries was performed in all subjects. The primary study outcome measurements were histopathologic plaque features determining severity of atherosclerosis, including %stenosis, calcification, thin-cap fibroatheromas (TCFA), and thrombotic CAD.

**Results:** After exclusion of cases with insufficient DNA sample quality or with missing data, 954 cases (mean age 48.8±14.7; 75.7% men) remained in the final study cohort. Subjects in the highest PRS quintile exhibited more severe atherosclerosis compared to subjects in the lowest quintile, with greater %stenosis (80.3%±27.0% vs. 50.4%±38.7%; adjusted p<0.001) and a higher frequency of calcification (69.6% vs. 35.8%; adjusted p=0.004) and TCFAs (26.7% vs. 9.5%; adjusted p=0.007). Even after adjustment for traditional CAD risk factors subjects within the highest PRS quintile had higher odds of severe atherosclerosis (i.e., ≥75% stenosis; adjusted OR 3.77; 95%CI 2.10-6.78; p<0.001) and plaque rupture (adjusted OR 4.05; 95%CI 2.26-7.24; p<0.001). Moreover, subjects within the highest quintile had higher odds of CAD-associated cause of death, especially among those aged 50 years and younger (adjusted OR 4.08; 95%CI 2.01-8.30; p<0.001). No associations were observed with plaque erosion.

**Conclusions:** This is the first autopsy study investigating associations between PRS and atherosclerosis severity at the histopathologic level in subjects with sudden death. Our pathological analysis suggests PRS correlates with plaque burden and features of advanced atherosclerosis and may be useful as a method for CAD risk stratification, especially in younger subjects.

**Highlights:**

- In this autopsy study including 954 subjects within the CVPath Sudden Death Registry, high PRS correlated with plaque burden and atherosclerosis severity.
- The PRS showed differential associations with plaque rupture and plaque erosion, suggesting different etiologies to these two causes of thrombotic CAD.
- PRS may be useful for risk stratification, particularly in the young. Further examination of individual risk loci and their association with plaque morphology may help understand molecular mechanisms of atherosclerosis, potentially revealing new therapy targets of CAD.

**Graphic Abstract:** A polygenic risk score, generated from 291 known CAD risk loci, was assessed in 954 subjects within the CVPath Sudden Death Registry. Histopathologic examination of the coronary arteries was performed in all subjects. Subjects in the highest PRS quintile exhibited more severe atherosclerosis as compared to subjects in the lowest quintile, with a greater plaque burden, more calcification, and a higher frequency of plaque rupture.

## Introduction

Cardiovascular diseases (CVD) remain the leading causes of death worldwide.^1^ Despite universal use of risk stratification algorithms based upon traditional cardiovascular risk factors (CVRF) and preventative therapies such as lipid lowering, significant residual risk for CVD remains. While cardio-metabolic, behavioral, environmental, and social factors are key drivers for CVD, genetic factors are estimated to account for 30–60% of the variation in risk.^2–4^ Over the last decade, large-scale genome-wide association studies (GWAS) have identified genetic variants associated with an elevated risk for coronary artery disease (CAD).^5–12^ As of 2022, 328 common genetic loci associated with CAD at a genome-wide significance level have been described.^13–15^

Genetic risk determination might potentially allow for an earlier identification of individuals at high risk for CAD. Polygenic risk scores (PRS), derived as the weighted sum of the total number of risk alleles in an individual, have been used to quantify the genetic risk for CAD in large prospective cohorts, and higher PRS correlated with cardiovascular events.^16–18^ Moreover, subjects with high PRS may derive larger benefit from healthy lifestyle^18^ and cholesterol-lowering treatment^17^ compared with those with low PRS. Because PRS can be assessed at birth, they offer the possibility to identify individuals at high risk for atherosclerosis development long before the onset of CVRF or CAD. Furthermore, a recent study investigating associations between PRS and plasma protein levels in CVD showed PRS can be used to identify proteins through which detrimental effects arise and which may serve as potentially druggable targets.^19^

PRS for CAD are typically constructed based on genetic findings of large-scale CAD GWAS. However, as most GWASs have been performed in populations of European ancestry^14, 20–23^, and large-scale GWAS of CAD have been conducted only recently in multiple ancestral groups^15, 24–26^, variants associated with risk are disproportionately common and discovered in European populations, making it difficult to understand to what extent PRS are transferable to groups of different ancestries.^27^

Furthermore, it has never been studied whether PRS associate with autopsy findings describing the various atherosclerosis morphologies that characterize CAD. For instance, plaque rupture and erosion are two morphologic entities underlying acute myocardial infarction (MI) which likely have distinct pathogeneses. Studying associations between PRS and specific plaque phenotypes might serve as a basis for exploring the underlying molecular pathways and mechanisms of increased cardiovascular risk. A better understanding of the genetic factors that predict risk for each might offer new insights into how to prevent atherosclerosis and aid in the development of new therapeutic targets.

We sought to determine for the first time whether previously identified genetic predictors of CAD associate with coronary atherosclerosis at autopsy. Based on known CAD risk loci, we calculated PRS for 954 subjects in the CVPath Sudden Death Registry. Our primary hypothesis was that higher PRS might be associated with more severe atherosclerosis at the histological level. Atherosclerosis severity was hereby defined by the degree of atherosclerotic luminal narrowing of the coronary arteries as well as by histopathologic plaque features indicating plaque vulnerability, including calcification and the presence of a thin fibrous cap. Our secondary hypothesis was that higher PRS might also be associated with a higher frequency of coronary thrombosis, either triggered by plaque rupture or by plaque erosion. Finally, we hypothesized that higher PRS might consequently associate with CAD-related causes of death as certified by the autopsy report.

## Methods

A detailed description of all methods is provided in the **Supplemental Material.**^7, 9–11, 14, 15, 17, 18, 24–26, 28–40^

### Study population

The CVPath Sudden Death Registry comprises a total of 4,327 hearts from subjects ≥18 years who died of unexpected sudden death. Of these, 2,455 subjects were randomly selected for genotyping. Genotyping was performed in two separate datasets at two locations. The first dataset comprised 1,490 samples from self-reportedly White subjects, and genotyping was performed at Erasmus University of Rotterdam (“Erasmus dataset”). The second dataset comprised 965 samples from self-reportedly Black subjects, and genotyping was performed at the University of Texas UTHealth Human Genetics Center Laboratory (“SCA dataset”). 1,238 subjects were excluded because of insufficient DNA sample quality (for details, see **Supplemental Material**). In addition, 263 subjects were removed for missing or inconsistent data or because they did not match the study inclusion criteria (**Figure 1**). Baseline characteristics were not significantly different between subjects who were excluded and subjects who remained in the final analysis (**Supplemental Table S2**). The final study cohort comprised 954 sudden death cases. The protocol for the study was approved by the Institutional Review Board of CVPath Institute (Study ID: RP#0115), and a waiver of consent was granted. AC, NG, RV and AF had full access to all data and take responsibility for its integrity and the data analysis.

**Figure 1.**
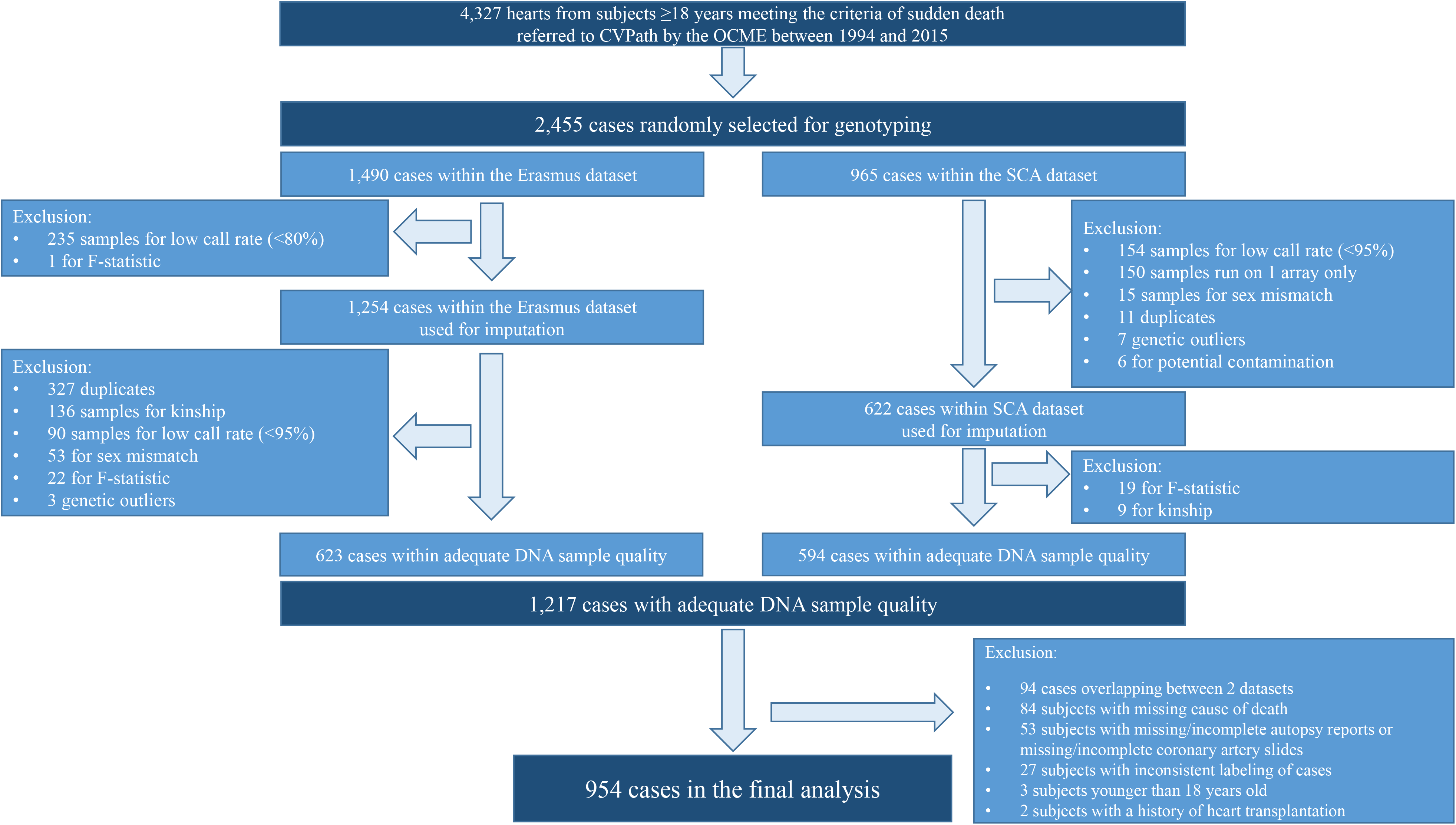
Study Cohort. From 4,327 subjects ≥18 years referred to CVPath between 1994 and 2015, 2,455 cases were randomly selected for genotyping. Genotyping was done in two datasets, the “Erasmus dataset”, comprising DNA from self-reportedly White subjects, and the “SCA dataset”, including DNA from self-reportedly Black subjects. After exclusion of 236 cases from the Erasmus dataset for quality control issues, data from 1,254 cases were used for imputation. 343 samples were removed from the SCA dataset before imputation. Additional QC on the imputed datasets removed 631 samples from the Erasmus dataset and 28 samples from the SCA dataset, leaving 1,217 cases with adequate sample quality. 94 cases overlapping between Erasmus and SCA were removed. 263 subjects were excluded for missing cause of death, incomplete autopsy reports, missing/incomplete coronary artery slides, inconsistent labeling, age younger than 18 years, or for a history of heart transplantation. The final study cohort comprised 954 cases.

### PRS construction

From 328 known common CAD risk loci^12^, 291 loci were available for PRS calculation (**Supplemental Table S3**). PRS were computed using PRSice^41^ by summing the number of risk alleles across all loci, weighted by the effect size of that SNP from previous GWAS. For details, see **Supplemental Material**. We sought to find associations between PRS and coronary atherosclerotic plaque morphologies, including calcification, the presence of thin-cap fibroatheromas (TCFAs), and the degree of atherosclerotic narrowing of the coronary arteries. Furthermore, we tested associations between PRS and coronary thrombosis, including plaque rupture and plaque erosion. Finally, we evaluated whether PRS was associated with CAD-related causes of death.

### Statistics

All analyses were first performed in a dataset-stratified fashion with ancestry-specific construction of principal components (PCs) and separate standardization of the PRS using z-scores in the SCA and in the Erasmus datasets. In addition, a pooled analysis was performed using PCs generated for the pooled sample and additional adjustment for self-reported race. In each cohort, subjects were divided into quintiles according to the standardized PRS. Logistic regression analyses were performed to compare the odds of binary outcomes among PRS quintiles. The first quintile served as reference. Model 1 was adjusted for age, self-reported race, sex, and the first 10 PCs, and model 2 was additionally adjusted for BMI and traditional CVRF (*i.e.*, hypertension, smoking, diabetes mellitus, and hyperlipidemia); for details see **Supplemental Material**. Generalized linear mixed model analysis was performed to determine associations between PRS and continuous variables with adjustment for the above-mentioned covariates and using the study identifier of each subject as random effect variable. Multiple testing correction was performed using Tukey’s multiple comparisons test where applicable. Statistical analyses were performed with SPSS version 28.0. Continuous data are presented as means ± SD. Categorical data are presented as numbers and percentages. Graphs were created in GraphPad Prism (GraphPad Software, Version 8.4.1).

## Results

### Overall characteristics

Baseline characteristics of 562 self-reportedly Black subjects within the SCA dataset and 392 self-reportedly White subjects within the Erasmus dataset with available genomic data are summarized in **Supplemental Table S4**. Self-reportedly Black subjects died at significantly younger age compared to self-reportedly White subjects within with a mean age gap of approximately 7.4 years. Furthermore, the percentage of women was higher among self-reportedly Black subjects. Self-reportedly white subjects had more severe atherosclerosis, as indicated by greater cross-sectional luminal narrowing and more advanced plaque features, including calcification and TCFAs. Furthermore, plaque rupture was more frequent in self-reportedly White than in Black subjects.

The SCA and the Erasmus cohorts were divided into quintiles according to z-score-standardized PRS. There were no differences in CVRF between the lowest (the 1^st^) and the highest (the 5^th^) PRS quintiles for either group (**Table 1**). In the Erasmus dataset, subjects within the fifth quintile died at significantly younger age compared to subjects within the first quintile (52.0 ± 13.9 years vs. 57.4 ± 17.2 years).

**Table 1.**
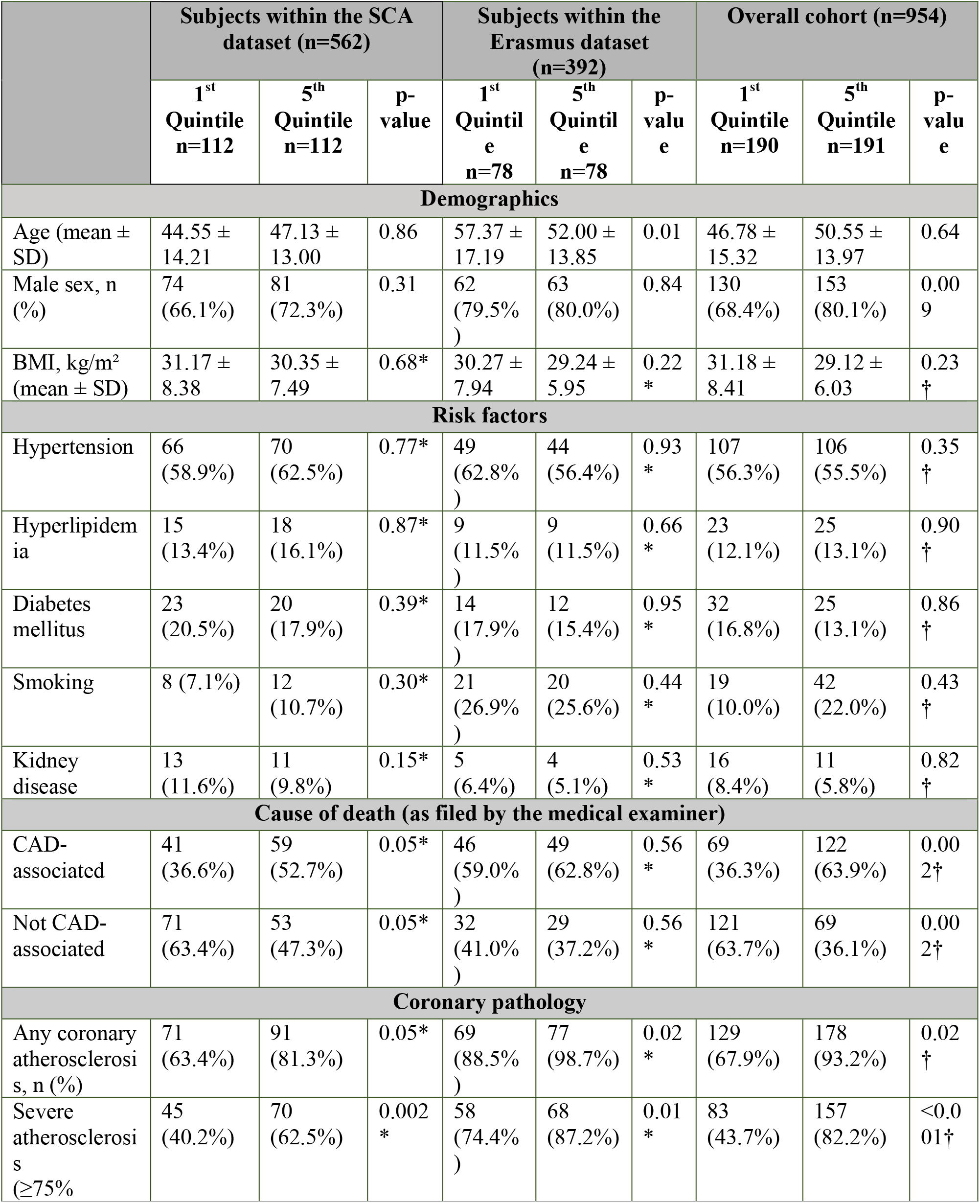
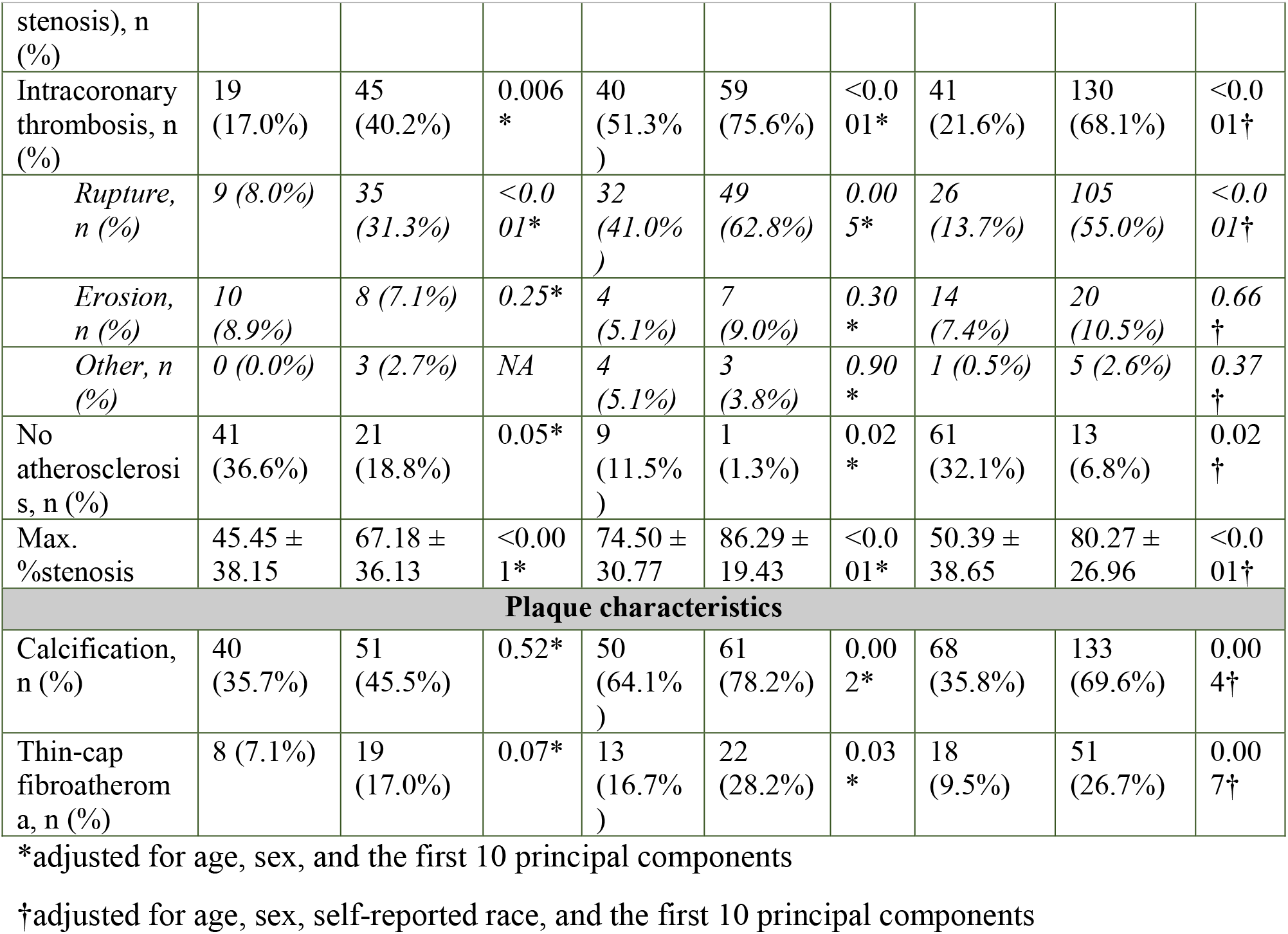
Comparisons Between the Highest and the Lowest PRS Quintiles

### Associations with luminal narrowing

Detailed histopathologic examination of the coronary arteries was performed in each subject. The maximum percentage of cross-sectional luminal narrowing (*i.e.*, max. %stenosis) was on average significantly higher in subjects within the race-specific highest PRS quintile compared to the lowest quintile after accounting for the impact of age, sex, self-reported race, and the first 10 PCs (80.3% vs. 50.4%, adjusted p<0.001 in the overall analysis; 67.2% vs. 45.5%, adjusted p<0.001 in subjects within SCA; 86.3% vs. 74.5%, adjusted p<0.001 in subjects within Erasmus; **Table 1**, **Figure 2**). This was still true when the analysis was stratified by the proximal, middle, or distal part of the coronary arteries and when each coronary artery was considered separately (**Supplemental Figure S3**). After additional adjustment for traditional CVRF (**Supplemental Table S5**), max. %stenosis was 18.4% higher in the highest compared to the lowest quintile in the overall cohort (estimated marginal means 77.8% vs. 59.4%; adjusted p<0.001). 16.7% higher max. %stenosis in subjects within the highest compared to the lowest quintile was observed in the SCA dataset (71.5% vs. 54.8%; adjusted p<0.001), and in Erasmus, subjects within the highest quintile had 16.4% higher max. %stenosis compared to the lowest quintile (82.0% vs. 65.6%; adjusted p<0.001).

**Figure 2.**
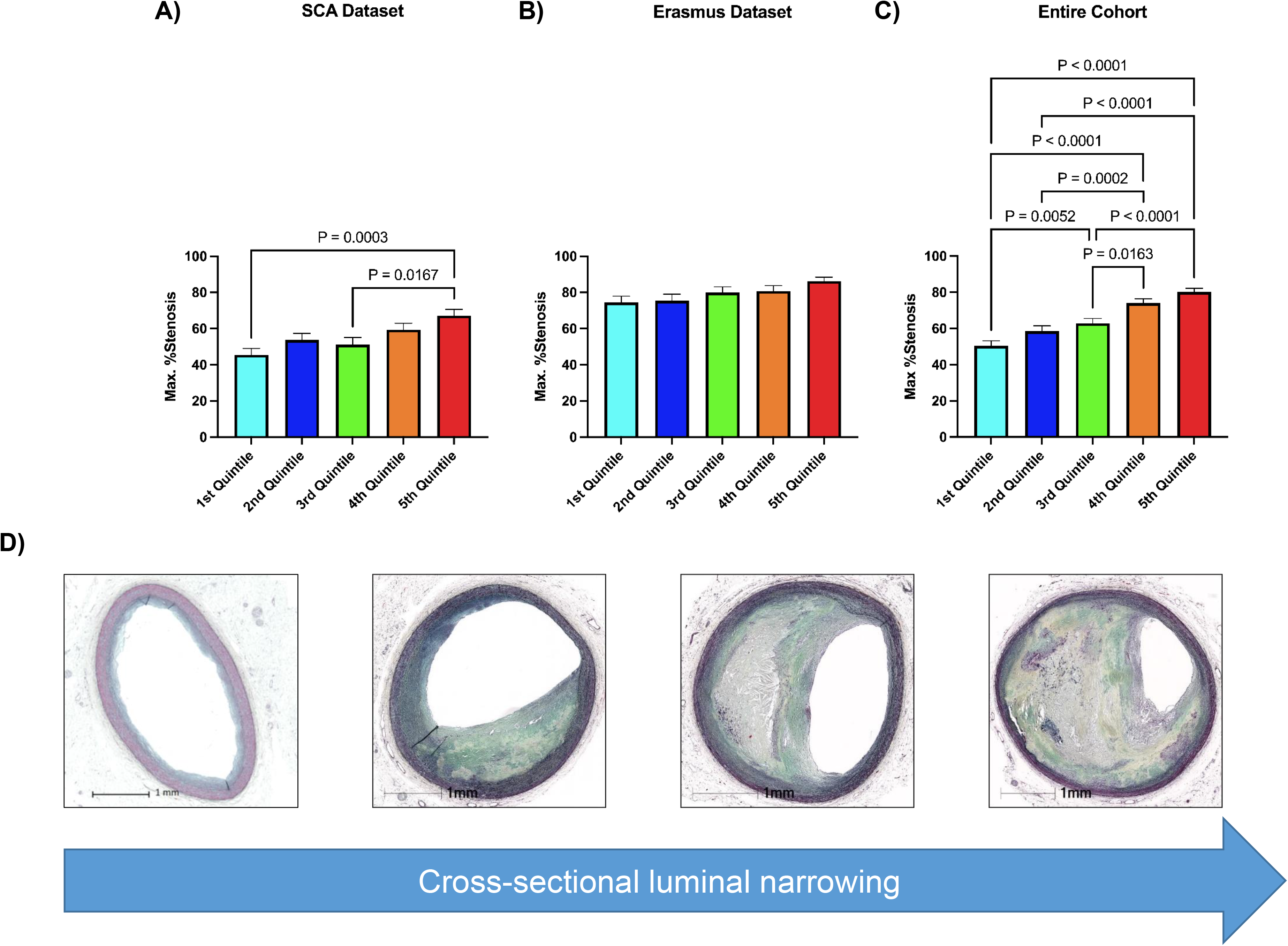
Associations between PRS and Maximum %Stenosis. A-C) Subjects within the highest quintile of standardized PRS exhibited a higher maximum percentage of cross-sectional luminal narrowing (*i.e.*, max. %stenosis) in histology compared to subjects within the lowest PRS quintile, with statistical significance both in the SCA dataset **(A)** and in the combined analysis **(C)**. **D)** Representative histopathologic images of coronary arteries with increasing %stenosis (from left to right: no /negligible atherosclerosis (18.7% stenosis), mild (46.9% stenosis), moderate (67.5% stenosis), and severe atherosclerosis (85.9% stenosis)).

To evaluate if the observation of higher %stenosis in higher PRS quintiles was age-dependent, we divided our cohort in two groups of subjects under the age of 50 (n=595) and subjects of 50 years and older (n=359). In subjects under the age of 50 years, we observed a stepwise increase in max. %stenosis with higher quintiles, with a statistically significant difference in %stenosis between the 1^st^ and the 5^th^ quintile in all three datasets (**Supplemental Figure S4, A-C; Supplemental Table S6**). In subjects of 50 years and older, only in the overall cohort subjects within the 5^th^ quintile had higher max. %stenosis compared to the 1^st^ quintile while there were no significant differences in %stenosis when the SCA and the Erasmus datasets were considered separately (**Supplemental Figure S4, D-F**).

Subjects with severe atherosclerosis (*i.e.*, ≥75% cross-sectional stenosis in histology; n=591) ranked higher in z-score-standardized PRS and PRS percentiles compared to those with no severe atherosclerosis (n=363; **Supplemental Figure S5, A-F**). The finding of severe atherosclerosis was more likely among subjects in the highest compared to the lowest PRS quintile, both after adjustment for age, sex, self-reported race, and the first 10 PCs in model 1 (**Figure 3**) and after additional adjustment for traditional CVRF in model 2 (**Figure 4; Supplemental Figure S5G**).

**Figure 3.**
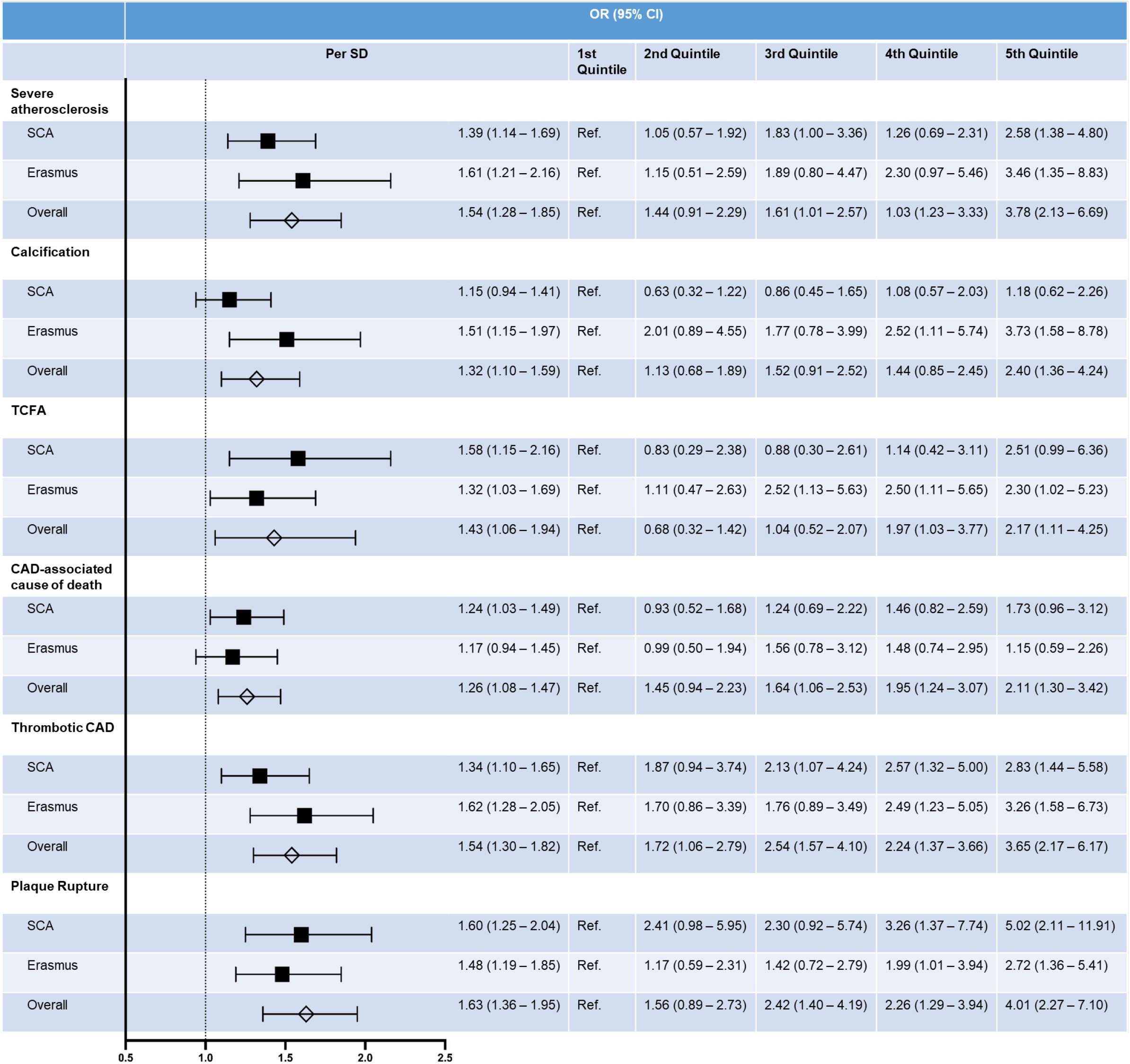
Associations between PRS and Atherosclerotic Endpoints in Model 1. Odds ratios for histopathologic features of atherosclerosis including severe atherosclerosis (defined as ≥75% luminal narrowing), calcification and TCFA, as well as for CAD-associated cause of death, thrombotic CAD and plaque rupture per each standard deviation increase and according to PRS quintiles, adjusted for age, sex, and the first 10 race-specific PCs (model 1).

**Figure 4.**
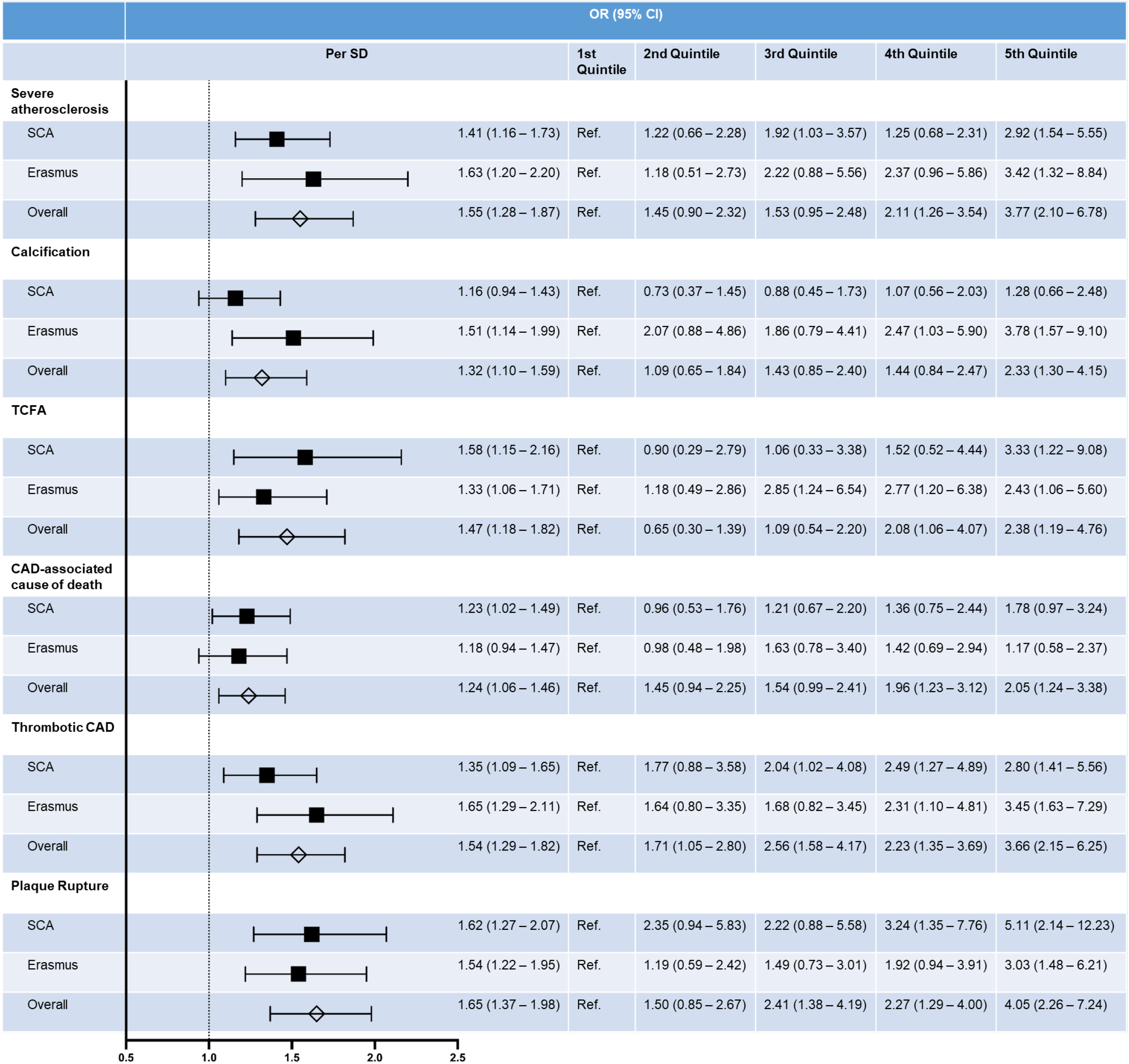
Associations between PRS and Atherosclerotic Endpoints in Model 2. Odds ratios for atherosclerotic endpoints per each standard deviation increase and according to PRS quintiles with adjustment for age, sex, BMI, hypertension, hyperlipidemia, diabetes, smoking, and the first 10 race-specific PCs (model 2).

To further explore whether PRS was associated with the extent of clinically relevant CAD, we compared PRS among subjects with severe three-, two-, one- or “zero”-vessel disease. Subjects with severe three-vessel disease were defined as having ≥75% cross-sectional luminal narrowing of all three major epicardial coronary arteries (*i.e.*, LAD, LCX, and RCA) in histology, whereas subjects with “zero”-vessel disease had no severely narrowed artery (*i.e.*, the max. %stenosis was less than 75% in all arteries). Z-score standardized PRS was highest in subjects with three-vessel disease and lowest in subjects without severe CAD, and subjects with severe three-vessel disease ranged higher in PRS percentiles compared to subjects without severe CAD in all three cohorts (**Supplemental Figure S6**).

### Associations with advanced plaque features

Next, we evaluated whether higher PRS was associated with features of advanced atherosclerosis. In all three datasets, subjects with coronary artery calcification ranked higher in z-score standardized PRS compared to subjects without calcified plaques (**Supplemental Figure S7**). After adjustment for covariates in model 1, coronary artery calcification was 2.4 times more likely in the highest compared to the lowest PRS quintile in the overall cohort, and 3.7 times more likely in the highest compared to the lowest quintile in the Erasmus dataset (**Figure 3**). Higher odds of coronary artery calcification in the highest compared to the lowest PRS quintiles were still observed in the overall cohort and in the Erasmus dataset after additional adjustment for BMI and traditional CVRF in model 2 (**Figure 4**). In the SCA dataset, however, the odds for calcification were not significantly higher in the highest compared to the lowest PRS quintile after adjustment for covariates (**Figures 3, 4**).

Subjects with histologic finding of TCFAs ranked higher in z-score standardized PRS compared with subjects without TCFAs (**Supplemental Figure S8**). The frequency of TCFAs was higher in coronary arteries from subjects within the highest compared with the lowest PRS quintile (**Table 1**), and higher odds of TCFA were still present after adjustment for covariates in models 1 and 2 (**Figures 3, 4**).

### Associations with coronary thrombosis

Coronary thrombosis was defined as intraluminal coronary thrombus formation, either from plaque rupture or plaque erosion, or from other reasons (e.g., calcified nodules). Subjects with histologic evidence of coronary thrombosis ranked higher in PRS compared to those without thrombosis (**Supplemental Figure S9**). The finding of coronary thrombosis was more frequent in subjects within the highest compared with the lowest PRS quintile (**Table 1**), and coronary thrombosis was still associated with higher PRS after adjustment for covariates in both models (**Figures 3, 4**).

We explored whether PRS was different depending on the occlusion site (proximal vs. middle vs. distal parts of the coronary artery) and found no difference in PRS percentiles among subjects with occlusions in the proximal, middle, and distal part of the coronary arteries (**Supplemental Figure S10 A-C**). Likewise, PRS was not different in occlusions of the LAD vs. the LCX vs. the RCA (**Supplemental Figure S10 D-F**).

To further investigate whether higher PRS associated with specific causes of coronary thrombosis, we compared PRS in subjects with plaque rupture and plaque erosion to subjects with different stages of non-thrombotic CAD. Subjects with plaque rupture ranged highest in PRS percentiles in all three datasets. Subjects with plaque erosion, however, only ranged higher in PRS percentiles compared to subjects with no or only mild CAD in the overall cohort (**Supplemental Figure S11 A-C**). To evaluate whether there was a statistically significant difference in the odds for plaque erosion among PRS quintiles, we performed logistic regression in a subgroup including 95 subjects with erosion and 193 subjects with no CAD in order to avoid bias by subjects with more severe CAD and rupture in the comparison group. After adjustment for covariates in model 1, subjects in the highest PRS quintile had 3.9 times higher odds of plaque erosion compared to subjects in the lowest quintile. Significance was lost, however, in model 2 with further adjustment for BMI and traditional CVRF (**Supplemental Figure S11D**).

We observed a higher frequency of plaque rupture among subjects within the highest compared to the lowest PRS quintile in all three datasets (**Table 1**). Subjects with rupture also ranked higher in PRS percentile-wise compared to those without rupture (**Supplemental Figure S12**). Higher PRS was still associated with the finding of plaque rupture after adjustment for covariates in model 1 and in model 2 **(Figures 3, 4)**. Since plaque burden is a strong risk factor for plaque rupture, we performed additional adjustment for %stenosis in logistic regression. Indeed, higher %stenosis was strongly and independently associated with higher odds for plaque rupture after adjustment for covariates (OR 1.08; 95%CI 1.06-1.09 in the overall cohort; OR 1.07; 95%CI 1.05-1.10 in Erasmus; OR 3.13; 95%CI 1.16-8.45 in SCA). Nevertheless, in the SCA dataset and in the overall cohort, subjects within the fifth quintile of PRS still had higher odds for plaque rupture compared to subjects within the first quintile (**Supplemental Figure S12H**).

### Associations with CAD-related causes of death

We also investigated whether higher PRS was associated with a higher likelihood of dying from CAD-associated causes, as filed by the State of Maryland Medical Examiner. In the overall cohort, CAD-associated causes were more frequently filed in subjects within the highest PRS quintile compared to those within the lowest quintile. While there was a trend towards higher frequency of CAD-associated cause of death in the highest vs. the lowest PRS quintile in SCA, no significant difference was observed among subjects within Erasmus (**Table 1**). To further examine whether higher PRS might predispose to premature death from CAD, we separated our cohorts by age, with a cutoff set at 50 years. 286 out of 595 younger subjects (48.1%) and 207 out of 359 older subjects (57.7%) died from CAD-associated causes.

In all three datasets, younger subjects who died from a CAD-associated cause of death ranked higher in PRS percentiles compared to younger subjects who died from non-CAD-associated causes (**Supplemental Figure S13A**). After adjusting for covariates, higher odds of dying from CAD-associated causes with higher PRS remained (**Supplemental Figure S13B**), and younger subjects within the 1^st^ quintile were at approximately 4.1-times higher odds of dying from CAD-associated causes compared to younger subjects within the 5^th^ quintile (**Supplemental Figure S13C**).

Among subjects of 50 years and older, no difference in PRS was observed between subjects with CAD-associated causes of death and subjects who died of non-CAD-associated causes (**Supplemental Figure S13D**). Higher PRS was not associated with higher odds of CAD-associated causes of death after adjustment for covariates in logistic regression (**Supplemental Figure S13E, F**).

## Discussion

This is the first autopsy study investigating associations between a CAD GWAS-derived PRS and histopathologic findings of coronary atherosclerosis using a uniquely large cohort of human coronary arteries. While acknowledging that our final cohort was relatively small compared to PRS studies using clinical endpoints, our study provides information about the relationship between PRS to human atherosclerotic plaque characteristics on a level never performed previously.

### Significance of high PRS in CAD severity

In line with clinical studies, we found significant associations between high PRS, derived from 291 known genetic risk loci for CAD, and a higher degree of luminal narrowing, even after controlling for confounding variables. A study by Natarajan et al. including 1,154 participants in CARDIA (Coronary Artery Risk Development in Young Adults) showed associations between a PRS, derived from 57 common variants, and coronary artery plaque burden, assessed by coronary computed tomography angiography (CTA).^42^ Likewise, Christiansen et al. demonstrated higher segment stenosis scores as well as a higher plaque prevalence in 1,645 patients with suspected CAD undergoing coronary CTA.^43^ Furthermore, subjects with histopathological evidence of severe multivessel disease ranked higher in PRS percentiles compared to those without multivessel disease. Indeed, Hindieh et al. reported strong associations between a 30-variant PRS and the presence of multivessel disease in a cohort of 763 patients with premature acute coronary syndrome.^44^

Higher PRS was also associated with the presence of severe plaque features in our study, including calcification and TCFA. Indeed, numerous clinical studies have reported positive associations between high PRS and coronary artery calcification.^18, 42, 43, 45^ However, while calcification is relatively easy to assess and to quantify using coronary CTA, noninvasive identification of TCFAs can be challenging, and to the best of our knowledge, there is no clinical study investigating associations between genetic risk and the presence of TCFAs in the coronary tree. Results from clinical studies investigating associations between specific plaque features and PRS have been contradictory. While data from the Prospective Multicenter Imaging Study for Evaluation of Chest Pain (PROMISE) trial suggested a high proportion of high-risk CAD, defined as obstructive CAD, coronary artery calcification score >400, high extent of stenosis and plaque burden, or high-risk plaque features, including signs of positive remodeling, low attenuation, or napkin ring sign on CTA, in the highest PRS quintile in a cohort of 605 participants^46^, Christiansen et al. did not find associations between PRS and any specific plaque characteristics over others.^43^ In contrast to clinical studies, however, histopathology allows for a more detailed investigation of coronary artery plaque features which might have been missed in CTA or on the angiogram.

### PRS and histopathologic findings underlying MI

PRS was also associated with the finding of coronary thrombosis, mainly triggered by plaque rupture, in all three datasets. Numerous clinical studies have shown associations between high genetic risk and an increased risk of MI.^42, 47, 48^ However, previous studies have also shown that only few loci are specifically associated with MI while the vast majority simply increases CAD risk in general.^36, 40, 49^ Coronary atherosclerosis is a prerequisite for most MI cases, and both factors are strongly interrelated. We found significant associations between higher PRS and plaque rupture, even after adjustment for %stenosis, suggesting that PRS also reflects other factors which might additionally increase the risk, such as plaque vulnerability and platelet reactivity.^50^ Furthermore, some MIs may occur silently. While we were able to differentiate between subjects with and without signs of MI in histology, silent MIs are hard to capture in clinical studies. Further studies should be conducted to dissect associations between specific SNPs and MI.

High PRS was associated with plaque rupture. On the other hand, no consistent associations were observed with plaque erosion, although subjects showing this pathology had a higher PRS compared to those with no or mild CAD. This might suggest there is genetic association with rupture while erosion may be caused by non-genetic factors or by genetic risk loci other than those that were included in our PRS calculation. Alternatively, this discrepancy could be based on potential biases inherent to the GWAS from which CAD risk loci were derived. Plaque rupture is responsible for approximately 65% of MIs, while only about 30–35% are caused by erosion.^28, 51^ Thus, an overrepresentation of plaque rupture could be expected in clinical registries and consortia. Therefore, it is not surprising that CAD risk loci derived from GWAS are mainly associated with rupture rather than with erosion. Likewise, the power to show associations with plaque erosion was lower in the present study. To fully answer the question whether genetic factors might have a role in the occurrence of plaque erosion, GWAS exclusively pertained to patients with erosion would be required. However, erosion remains a diagnosis of exclusion *in vivo* and can be confirmed unequivocally only in histopathology. Moreover, considering that a minority of MIs are caused by erosion, inclusion of a reasonable number of these types of events in order to achieve a sufficient sample size would presumably present a major challenge.

### PRS and coronary events in ancestrally diverse cohorts

Our study was conducted in two datasets; the Erasmus cohort including White subjects and the SCA dataset which consisted of Black Americans. Although there were some important differences in baseline characteristics between Black and White subjects, we found similarly strong associations between PRS and outcomes in both cohorts. Nevertheless, most GWAS and clinical studies investigating PRS have been performed in subjects of European ancestry^5, 52–54^ and it has been only recently that large-scale GWAS studies have been conducted in ancestrally diverse cohorts.^15^ Thus, more studies are warranted to uncover further genetic risk loci that might be specifically relevant in subjects from other ethnicities. However, our findings are in line with clinical studies. Khera et al. reported similar CAD risk prediction by PRS, constructed from 50 known CAD risk loci, among Black and White participants in ARIC.^18^ Ke et al. found that the majority of 71 CAD risk loci exhibited consistent directions in Black, Latino and Japanese participants in the Multiethnic Cohort^55^, and higher PRS constructed from these risk loci was more strongly associated with CAD in Black Americans and Latinos than it was in Japanese Americans, although effect sizes were smaller as compared with European datasets.^55^ More recently, Vassy et al. reported significant associations of higher PRS for CAD with incident MI and cardiovascular death in a multiancestry midlife- and older-age cohort including 79,151 participants in the Million Veterans Program.^48^ However, it is important to acknowledge that race is a rather poor way to group people, even though it is current practice in most studies investigating genetic impact on disease risk. Particularly for Black Americans, not only do proportions of African, European, and Amerindigenous ancestry vary by individual, but genetic distance between populations in Africa is larger than between other continental ancestries.^23, 56^ Furthermore, even within a group of people with similar ancestry, the accuracy of PRS can vary^27^, hence even though we adjusted for ancestry-specific PCs in each cohort and for PCs generated for the pooled sample, we probably did not fully account for the diversity within our study. In addition, the notion that plaque rupture occurred more often among self-reportedly White subjects might reflect selection bias. It has been reported previously that White subjects on average have a higher socio-economic status, often resulting in a better survival rate after SCD. Thus, in White subjects, there might be a predominance of cases with more advanced atherosclerosis.

### Clinical utility of PRS in risk prediction

PRS analysis can be carried out at any stage from birth and could potentially help identify subjects at increased risk for CAD at very young age, so that primary preventive measures could be individualized to those at highest genetic risk, perhaps even if they are not eligible for primary prevention as per traditional CVRF. Data from UK Biobank suggested PRS improved cardiovascular risk stratification especially early in life when later-life CVRF are still unknown, whereas by middle age the improvement in risk stratification attributed to PRS was marginal.^57^ Indeed, higher PRS was associated with CAD-associated causes of death only in those aged 50 years and younger, and we observed significantly increased coronary artery luminal narrowing with higher PRS quintiles in all three cohorts only in younger subjects. From a clinical perspective, given the strong impact of traditional CVRF later in life, it is indispensable to integrate non-genetic risk factors including age, sex, BMI, smoking behavior, hypertension, diabetes and hyperlipidemia into a predictive model and the increment in predictive accuracy of PRS versus a clinical risk score for CAD prediction has been reported to be modest ^58, 59^ or even absent.^60^ The positive predictive value of PRSs has been reported to be low, both in diagnostic settings and in prognosis estimation.^61^ It is unclear if higher PRS alone qualifies for a more intense screening for CAD, most likely in the context of guideline-recommended clinical risk assessment.^62^ Another obstacle is that the mere PRS value does not stand for itself but must always be interpreted in relation to population data. Still, PRSs may have great potential in research. It is likely that drivers of CAD are person- (not population) based, and specific targeted therapies based upon genetic predispositions may be needed in some subjects. While lipid-lowering therapies have been a mainstay of CAD prevention, other potential targets such as inflammation remain incompletely understood. Thus, mechanistic experiments are warranted to further explore whether or how individual or subgroups of SNPs included in the PRS might affect pathways that are known to be involved in atherogenesis. Further examination of morphological plaque features (*i.e.,* inflammation, lipid deposition, collagen content, etc.) and their association with subgroups of risk alleles may be an avenue towards a better understanding of the mechanisms by which genetics have impact on atherosclerosis, and potentially form a basis for the development of novel agents preventing the progression of CAD.

## Limitations

Besides limitations that have already been mentioned in earlier paragraphs, some additional limitations need to be highlighted. First, our dataset was quite select, with all subjects having died from suspected SCD, and therefore associations between PRS and CAD-associated causes of death need to be treated with caution. It is worth mentioning that samples have been collected since 1994 and the frequency of additional risk factors might have shifted over the years (e.g. smoking has diminished over the last years). In addition, the frequency of some CVRF, especially of smoking status and hyperlipidemia, might be underestimated, due to a lack of reporting in the autopsy reports.

We also need to emphasize once more that a majority of our samples did not withstand the rigorous QC criteria which is probably due to the nature of the study. First, in contrast to clinical genotyping studies, most samples were not obtained freshly. There is always a *post-mortem* interval to consider, including the time span between the subject’s death and finding the body. In our study, unexpected sudden death was defined as symptoms commencing within 6 hours of death (witnessed arrest) or death occurring within 24 hours after the victim was last seen alive in his normal state of health. Thus, the maximum time is limited to 24 hours, but nevertheless, depending humidity, temperature, and other environmental factors, we cannot exclude that this issue had impact on DNA quality. Second, DNA was extracted from either fresh frozen tissue (with all caveats to consider when using the word “fresh”) or formalin-fixed paraffin-embedded tissue blocks. Therefore, both the freezing and thawing processes and the formalin fixation procedure might limit the DNA quality. Nevertheless, we did not find significant differences in baseline characteristics between subjects who were excluded and those that were kept in our study cohorts. Despite the large portion of sample elimination due to strict inclusion criteria, this is still the largest heart cohort with both genetic and well-characterized histological data to date.

## Conclusion

This is the first autopsy study investigating associations between PRS, derived from 291 CAD risk alleles, and specific atherosclerotic plaque morphologies in coronary arteries. PRS was associated with cross-sectional luminal narrowing and features of advanced atherosclerosis, including calcification and the presence of TCFAs. Furthermore, PRS was associated with the occurrence of coronary thrombosis triggered by plaque rupture, while there were no consistent associations with plaque erosion. Highlighting PRS assessment might be of greater value in younger patients, higher PRS was positively associated with CAD-associated causes of death among subjects within the SCA dataset only in those aged 50 and younger. Further studies are warranted to investigate whether considering PRS for risk assessment in primary prevention will lead to improvements in clinical outcomes.

## Acknowledgements, Sources of Funding, & Disclosures

## a) Acknowledgements

We would like to acknowledge Ms. Laura Antonia Thiede, MS, and Ms. Nadiyeh Rouhi, MS, for their help with statistical analysis and figure design.

## Supplemental Material

Supplemental Methods

Supplemental Figures S1-S13

Supplemental Tables S1-S6

Major Resources Table

## Abbreviation List

CVD: cardiovascular disease
CVRF: cardiovascular risk factors
GWAS: genome-wide association studies
CAD: coronary artery disease
PRS: polygenic risk score
MI: myocardial infarction
TCFA: thin-cap fibroatheroma
PC: principal component
CTA: computed tomography angiography

